# Copy number variation on *ABCC2-DNMBP loci* impacts the diversity and composition of the gut microbiota in pigs

**DOI:** 10.1101/2022.10.06.510490

**Authors:** Yuliaxis Ramayo-Caldas, Daniel Crespo-Piazuelo, Jordi Morata, Olga González-Rodríguez, Cristina Sebastià, Anna Castello, Antoni Dalmau, Sebastian Ramos-Onsins, Konstantinos G. Alexiou, Josep M. Folch, Raquel Quintanilla, Maria Ballester

## Abstract

**Background:** Genetic variation in the pig genome partially modulates the composition of porcine gut microbial communities. Previous studies have been focused on the association between single nucleotide polymorphisms (SNPs) and the gut microbiota, but little is known about the relationship between structural variants and gut microbial traits.

**Results:** The main goal of this study was to assess the effect of porcine genome copy number variants (CNVs) on the diversity and composition of pig gut microbiota. For this purpose, we used whole-genome sequencing data to undertake a comprehensive identification of CNVs followed by a genome-wide association analysis between the estimated CNV status and the gut bacterial diversity in a commercial Duroc pig population. A CNV predicted as gain (DUP) partially harboring *ABCC2-DNMBP loci* was associated with richness (*p*-value=5.41×10^−5^) and Shannon α-diversity (*p*-value=1.42×10^−4^). The *in-silico* predicted gain of copies was validated by real-time quantitative PCR (qPCR), and its segregation, and positive association with the richness and Shannon α-diversity of the porcine gut bacterial ecosystem was confirmed in an unrelated F1 (Duroc×Iberian) cross. Furthermore, despite genetic and environmental differences between both populations, the gut microbiota of DUP samples showed a significant over-abundance of the *Desulfovibrio, Blautia, Phascolarctobacterium, Faecalibacterium, Succinivibrio* and *Anaerovibrio* genera.

**Conclusions:** In summary, this is the first study that evaluate the putative modulatory role of CNVs on pig gut microbiota. Our results advice the relevance of considering the role of host-genome structural variants as modulators of microbial ecosystems, and suggest the *ABCC2-DNMBP* CNV as a host-genetic factor for the modulation of the diversity and composition of the gut microbiota in pigs.

## BACKGROUND

Gut microbiomes have a profound impact on many aspects of pig health, such as the modulation of metabolic functions, physiological processes, and relevant porcine traits like growth [1], feed efficiency [2] [3], and immunocompetence [4]. Host-microbiome interactions are mediated by both environmental and host factors. Among them, genetic variation in the pig genome can modulate, in a taxa-specific manner, the composition and function of the pig gut eukaryotic and prokaryotic communities. Pig gut microbiota is heritable to an extent, showing low to medium heritabilities [2] [5] [6]. Quantitative trait loci (QTLs), genetic variants, and candidate genes associated to pig gut microbiota have been reported [7] [8] [9] [10].

However, since previous studies were focused on the association between SNPs and microbial traits, little is known about the relationship between the gut microbiota and structural variants in the porcine genome. Copy-number variants (CNVs) are structural variants that produce a change in the number of copies (gain or loss) of a genomic region. Compared to SNPs, CNVs involve large DNA segments that span a significant proportion of the genome, and account for greater genomic variability than SNPs. Consequently, CNVs are a relevant source of genetic variation that contribute to evolutionary adaptations, variation in gene expression and phenotypic traits in human and domestic animals [11] [12]. In humans, gain of copies of salivary amylase (*AMY1*) gene was associated with oral and gut microbiome composition [13]. In this seminal study, Poole et al. found that individuals with greater number of copies of *AMY1* showed greater levels of salivary *Porphyromonas*, followed by an increased abundance of resistant starch-degrading microbes in the gut.

We hypothesized that alike humans, CNVs are likely to contribute to animal gut microbial variability. However, to the best of our knowledge, such associations have not been documented in livestock. Consequently, the putative modulatory role of CNVs on the diversity, composition and function of livestock gastrointestinal microbiota remains to be elucidated. The main goal of this study was to assess the effect of porcine CNVs on the diversity and composition of pig gut microbiota.

## MATERIAL AND METHODS

### Animal samples

Samples employed in this study are a subset of pigs reported in [14] and [9]. In brief, a total of 100 weaned piglets (50 males and 50 females) from a commercial Duroc pig line were used as a discovery dataset (Table 1). The pigs were distributed in three batches, all animals were raised on the same farm and had *ad libitum* access to the same commercial cereal-based diet. Furthermore, a subset of 24 unrelated F1 (Duroc×Iberian) crossbred pigs with phenotypically extreme gut microbial diversity index (12 high and 12 low) from [7] were employed as independent validation dataset (Table 1).

**Table 1.** Descriptive statistics, mean and standard deviation (SD), of the richness and α-diversity in the discovery and validation datasets.

| Population | Dataset (n) | Groups | Richness (SD) | α-diversity (SD) |
| --- | --- | --- | --- | --- |
| Duroc (Discovery) | Complete (100) |  | 604.92 (98.17) | 6.05 (0.22) |
|  | Validate qPCR (48) | Diploid | 553.73 (92.71) | 5.95 (0.21) |
|  |  | DUP | 657.85 (89.70) | 6.16 (0.17) |
| Duroc x Iberic (Validation) | Complete (285) |  | 459.65 (104.33) | 5.82 (0.25) |
|  | Validate qPCR (24) | Diploid | 292.36 (67.18) | 5.36 (0.27) |
|  |  | DUP | 543.19 (158) | 6.06 (0.33) |

### Microbial DNA extraction, sequencing, and bioinformatics analysis

Fecal samples were collected from the Duroc piglets at 60□±□8 days of age and microbial DNA was extracted with the DNeasy PowerSoil Kit (QIAGEN, Hilden, Germany). Extracted DNA was sent to the University of Illinois Keck Center for paired-end (2□×□250 bp) sequencing on an Illumina NovaSeq (Illumina, San Diego, CA, USA). The 16S rRNA gene fragment was amplified using the primers V3_F357_N: 5’-CCTACGGGNGGCWGCAG-3’ and V4_R805: 5’-GACTACHVGGGTATCTAATCC-3’. Sequences were analysed with QIIME2 [15]; barcode sequences, primers, and low-quality reads (Phred score < 30) were removed. The quality control process also trimmed sequences based on expected amplicon length and removed chimeras. Afterwards, sequences were clustered into Amplicon Sequences Variants (ASVs) at 99% of identity. ASVs were classified to the lowest possible taxonomic level based on a primer-specific trained version of GreenGenes Database 13.8 [16]. Before the estimation of the diversity indices, to correct for the sequencing depth, samples were rarefied at 10,000 reads. Diversity metrics were estimated with the vegan R package v2.6-2 [17]. The α-diversity was evaluated with the Shannon index [18], and the β-diversity was assessed using the Whittaker index [19].

### Host-genome data analysis and CNV-calling

Simultaneously with fecal sampling, blood was collected at 60□±□8 days of age via the external jugular vein. Host genomic DNA was extracted from blood using the NucleoSpin Blood (Macherey–Nagel). Whole genome was paired-end sequenced (2 × 150 bp) in an Illumina NovaSeq6000 platform (Illumina) at *Centro Nacional de Análisis Genómico* (CNAG-CRG; Barcelona, Spain). Reads were mapped to the porcine reference assembly Sscrofa.11.1 with BWA-MEM 0.7.17 [20]. Alignment files containing only properly paired, uniquely mapping reads without duplicates were processed using Picard [http://broadinstitute.github.io/picard/] to add read groups and to remove duplicates. Variant calling was performed with the HaplotypeCaller tool from Genome Analysis Tool Kit (GATK 4.1.8.0) [21]. Applying GATK Best Practices, variants with minimum read depth of 5 on at least one sample were retained. Joint genotyping was conducted with combined gVCFs. Functional annotations were added using SnpEff v.5 [22] against the Sscrofa.11.1 reference database. CNV prediction was performed with ControlFREEC 11.5 [23], using a pool of samples as CNV baseline and using intervals of 20kb windows. CNV calls from all samples that were less than 10kb apart were merged with Survivor [24]. Individual CNV-calls were combined into copy number variant regions (CNVR) following the reciprocal overlap approach [11] with CNVRange [25]. Therefore, contiguous CNVs intervals with at least 50% of mutual overlap were merged into the same CNVR.

Nucleotide diversity pattern estimates were calculated considering the SNPs present in each individual separately. Here we tested two estimators: (i) Tajima’s theta estimator (π or nucleotide diversity, called ***ATajima*** hereafter), that is simply the number of variants present in the individual; (ii) ***RTajima*** estimator, which considers the frequency observed in the entire sample but calculated given the SNPs present in each individual. This estimate is equivalent to ***ATajima*** but needs to be corrected by the probability that only a portion of the total SNPs from a sample are present in each of the samples (by using a hypergeometrical distribution). Specifically, the calculation for a single individual is:

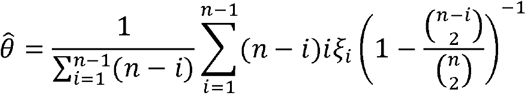

where *n* is the number of samples in the population (2×number of individuals) and *ξ_i_* is the number of SNPs observed in this individual that are at frequency *i* in the whole population.

The dataset in VCF format was converted to FASTA and from FASTA to transposed FASTA (tFASTA). This tFASTA file was read with the software *mstatspop* (https://github.com/CRAGENOMICA/mstatspop) to obtain the frequencies of SNPs from each of the pigs at the desired region. Finally, we calculated ***RTajima*** per fragment using self-made R scripts. All these estimates were finally divided by the effective length size of the studied region to obtain comparative estimates per nucleotide.

### CNV-wide association analysis

A genome-wide association analysis between the estimated CNV-status and gut bacterial diversity index was done using the following mixed model:

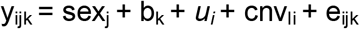

where y_ijk_ corresponds to the microbial index under scrutiny (richness or Shannon α-diversity) of the i-th individual animal of sex j in the k-th batch; sex_j_ and b_k_ correspond to the systematic effects of j-th sex (2 levels) and k-th batch (3 levels), respectively; *u*_i_ is the random additive genetic effect of the i-th individual, collectively distributed as 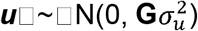 where 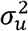 is the additive genetic variance and ***G*** is the numerator of the genomic relationship matrix calculated using the autosomal SNPs; cnv_li_ is the genotype (recoded as 11=loss, 12=diploid and 22=gain) for the l-th CNV of the i-th individual, and e_ijk_ is the residual.

### Quantitative real time PCR

Real-time quantitative PCR (qPCR) was used to validate the CNV on the *ABCC2* gene in a total of 72 samples including a subset of 48 Duroc samples (24 diploid and 24 *in silico* predicted as DUP), and 24 unrelated F1 Duroc×Iberian cross (Table 1). The CNV breakpoint was re-estimated with Manta: v1.6.0 [26]. All primers were designed using the Primer Express 2.0 software (Applied Biosystems). The pair of primers ABCC2_CNV_F 5’-*TGGCATCATTTATGTGGCTGTT-3’* and ABCC2_CNV_R 5’-*AGGAAGGAGCTTGGGCTTTTA-3’* amplify a specific region of the *ABCC2* gene containing the CNV, while the pair of primers ABCC2_F 5’-*TGGACAAGAACCAGAGTCAAAGC-3’* and ABCC2_R 5’-*ACATAGAGCGCATTTGAACGAA-3’* amplify a region outside of the estimated CNV breakpoint that was used as single copy control region. The 2-ΔΔCt method for relative quantification (RQ) of CNVs was used as previously described in [27]. qPCRs were carried out using SYBR Green chemistry (SYBRTM Select Master Mix, Applied Biosystems) and the instruments ABI PRISM® 7900HT and 7500 Real-Time PCR System (Applied Biosystems, Inc.; Foster City, CA). The reactions were carried out in a 20μl volume containing 10ng of genomic DNA. All primers were used at 900 nM. The thermal cycle was: 10 min at 95°C, 40 cycles of 15 sec at 95°C and 1 min at 60°C. Each sample was analyzed in quadrupled. PCR efficiencies (<95%) were evaluated with standard curves and dissociation curves were drawn for each primer pair to assess for the specificity of the PCR reactions. Three samples without CNV were used as reference. Results were analyzed with Thermo Fisher Cloud software 1.0 (Applied Biosystems), and qBase Plus v3.2 (Biogazelle).

### Identification of microbial signatures

The identification of ASVs that discriminate samples according to the number of copies of the *ABCC2-DNMBP loci* was performed based on the compositional kernel as implemented the function ‘*classify*’ of *kernInt* R package [28]. The ‘*classify*’ function run a supervised classification model based on Support Vector Machine. For that purpose, the available dataset was split at random into training set (80% of data) and validation set (20%). The *C* hyperparameter’s optimal value was obtained by 10×10 cross-validation on the training set. To estimate the mean classification accuracy the ‘classify’ function was run ten times using different training/test splits of the dataset. Microbial signatures were obtained from the hyperplane vector w, and the importance of the ASV k was computed by *kernInt* as (wk)^2^ [29]. Initially, the top 5% relevant taxa were retained, but a conservative approach was applied afterwards, keeping for subsequent analyses only the ASVs reported as relevant in at least the 50% of the replicates. Finally, to identify overrepresentation at genus level, the list of selected features was submitted to a taxa-set enrichment analysis [30].

## RESULTS

### Detection of copy number variants and association analysis

In this study we used whole-genome sequencing data from 100 healthy 60-day-old Duroc pigs to undertake a comprehensive identification of CNVs. A total of 1,292 CNVs distributed across 531 CNVR on autosomal pig chromosomes were identified (Figure 1, Supplementary table 1). After quality control, 1,005 CNVs grouped into 291 CNVR, presented in at least the 5% of the samples, were used for the association analysis. Among them, a CNV predicted as gain (DUP) located on CNVR454 (SSC14:111000000-111075999) that partially contain the ATP Binding Cassette Subfamily C Member 2 (*ABCC2*) and the Dynamin Binding Protein (*DNMBP*) genes showed a significant association with richness (p-value= 5.41×10^−5^) and the Shannon α-diversity (p-value=1.42×10^−4^) (Figure 2).

**Figure 1.**
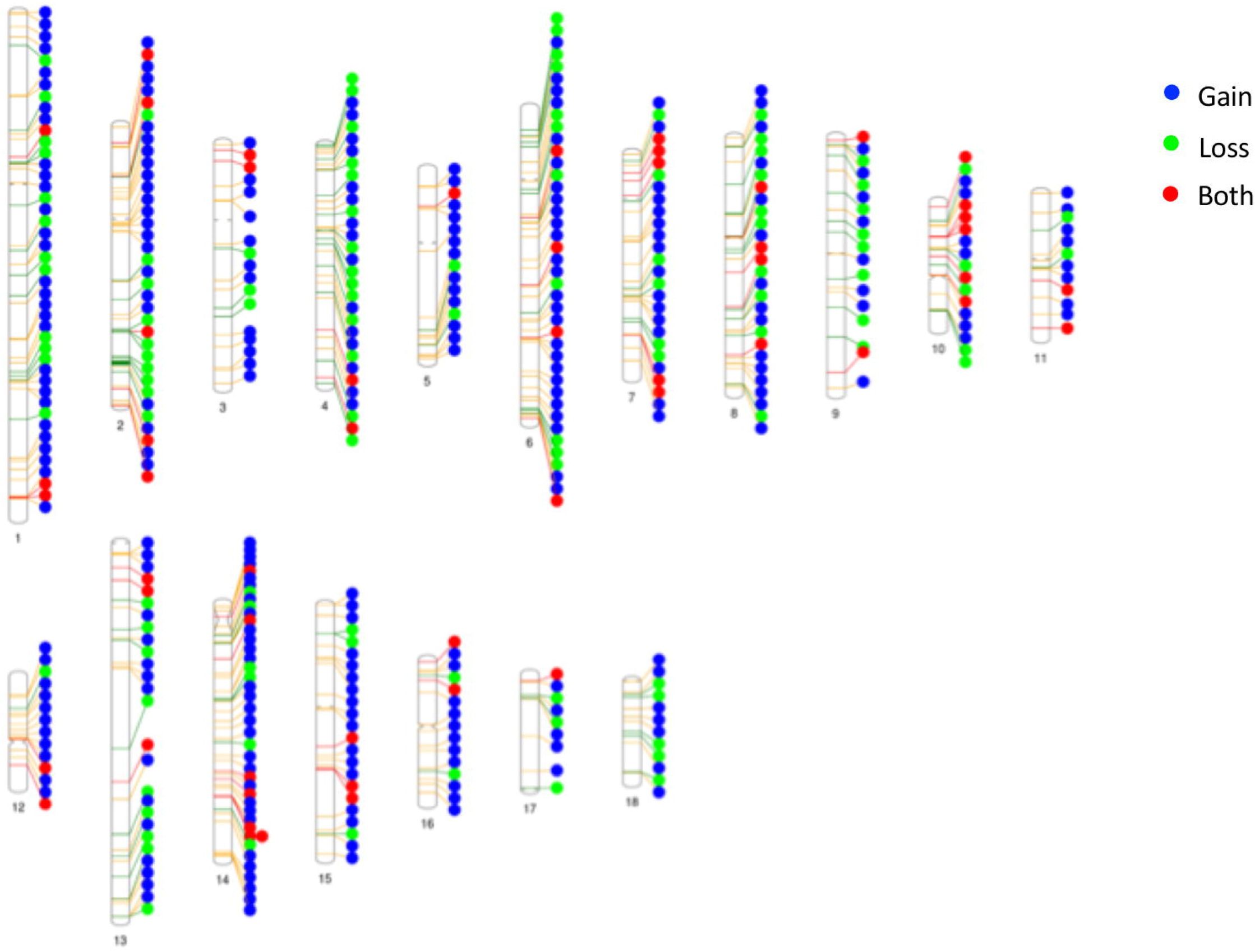
Graphical representation of the CNVRs detected. Green circles represent loss predicted status, gains are indicated in blue, and regions with either loss or gain status are represented in red. Chromosome sizes are represented in proportion to sequence length of the Sus scrofa 11.1 reference assembly.

**Figure 2.**
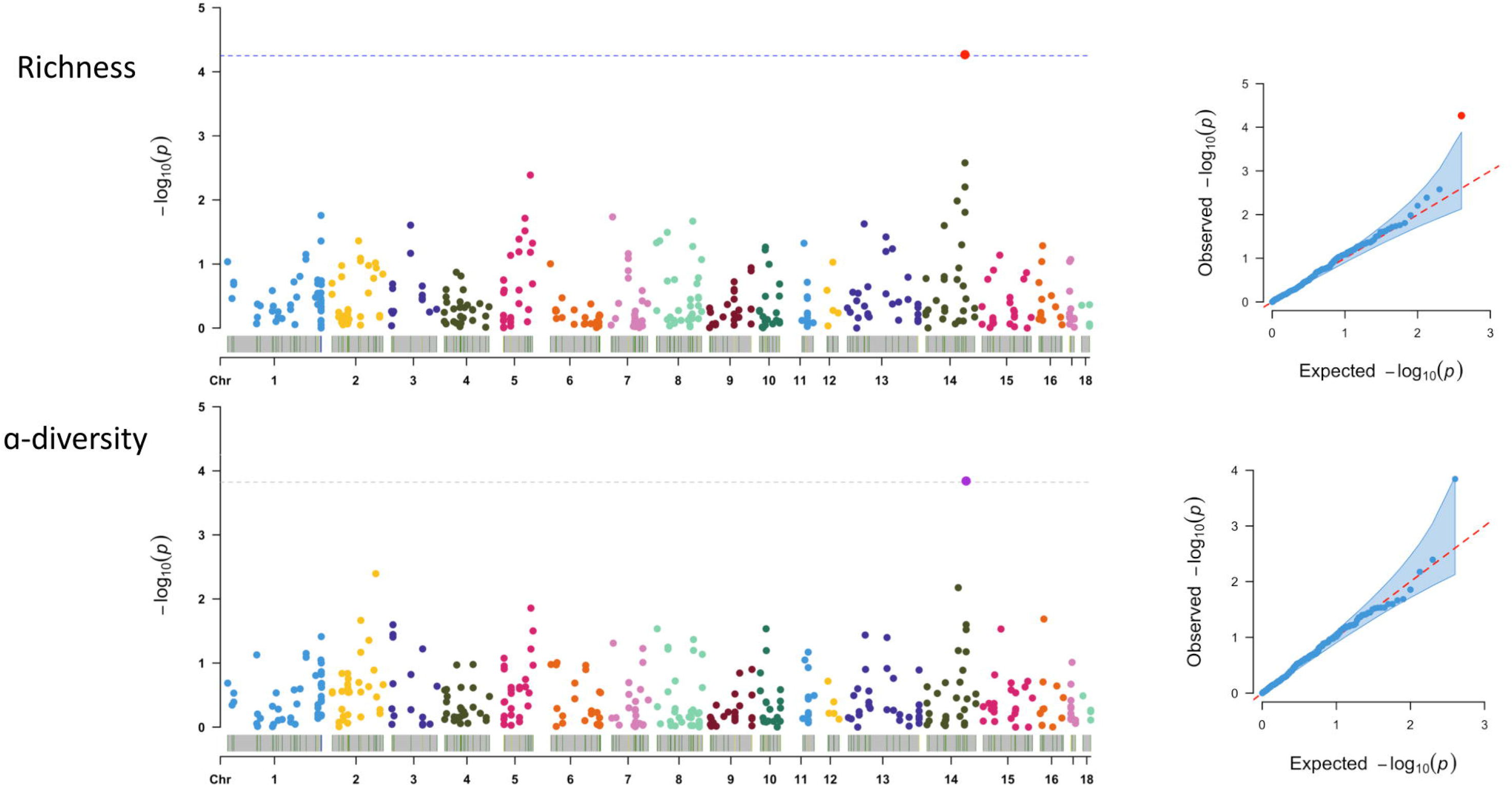
Results from the association analyses of CNVs identified across the pig genome with gut bacterial richness (A) and Shannon □-diversity (B). The x-axis represents the CNV position in the pig autosomal chromosomes (1–18), and the y-axis reflects the significance level represented as the −log10 (p-value).

### Validation by quantitative PCR

Since the *in silico* identification of CNVs may result in both false positive and negative results [27] [31], we conducted a qPCR assay with primers located on the *ABCC2* gene for the experimental validation of the CNV. The *in silico* predicted genotype was confirmed by qPCR in 46 out of the 48 Duroc samples, corresponding to an accuracy of 95.83%. To be noted, all the 24 samples *in silico* predicted as DUP presented the gain in number of copies. Thus, deviations from the diploid status were observed in two out of the 24 animals, where variation in the number of copies was not *in silico* predicted by ControlFREEC [23].

Because the existence of false negative sample assignation of CNV status could impact the GWAS results, and therefore, to avoid spurious associations and confirm our findings, we repeated the diversity index comparison using the subset of samples analyzed by qPCR (2N=22 vs DUP=26). In agreement with the CNV-GWAS, qPCR results corroborated that DUP pigs significantly had greater richness (p=1.8×10^−3^) and α-diversity values (p=3.8×10^−3^) (Table 1). Furthermore, RQ of the number of copies was positively correlated with the richness (r=0.474, p-value=6.72×10^−4^) and the α-diversity (r=0.401, p-value=7.77×10^−3^) (Figure 3). We also observed a positive relationship between the RQ of the number of copies and the nucleotide variability of the CNV genomic interval estimators ***ATajima*** (r=0.43) and ***RTajima*** (r=0.71). The high correlation observed between ***RTajima*** and the RQ values (Supplementary Fig 1) can be explained by the characteristic of RTajima statistic, which gives more importance to intermediate frequencies observed in the whole population, indicating that intermediate frequency diversity in this CNV is over-represented at greater number of copies.

**Figure 3.**
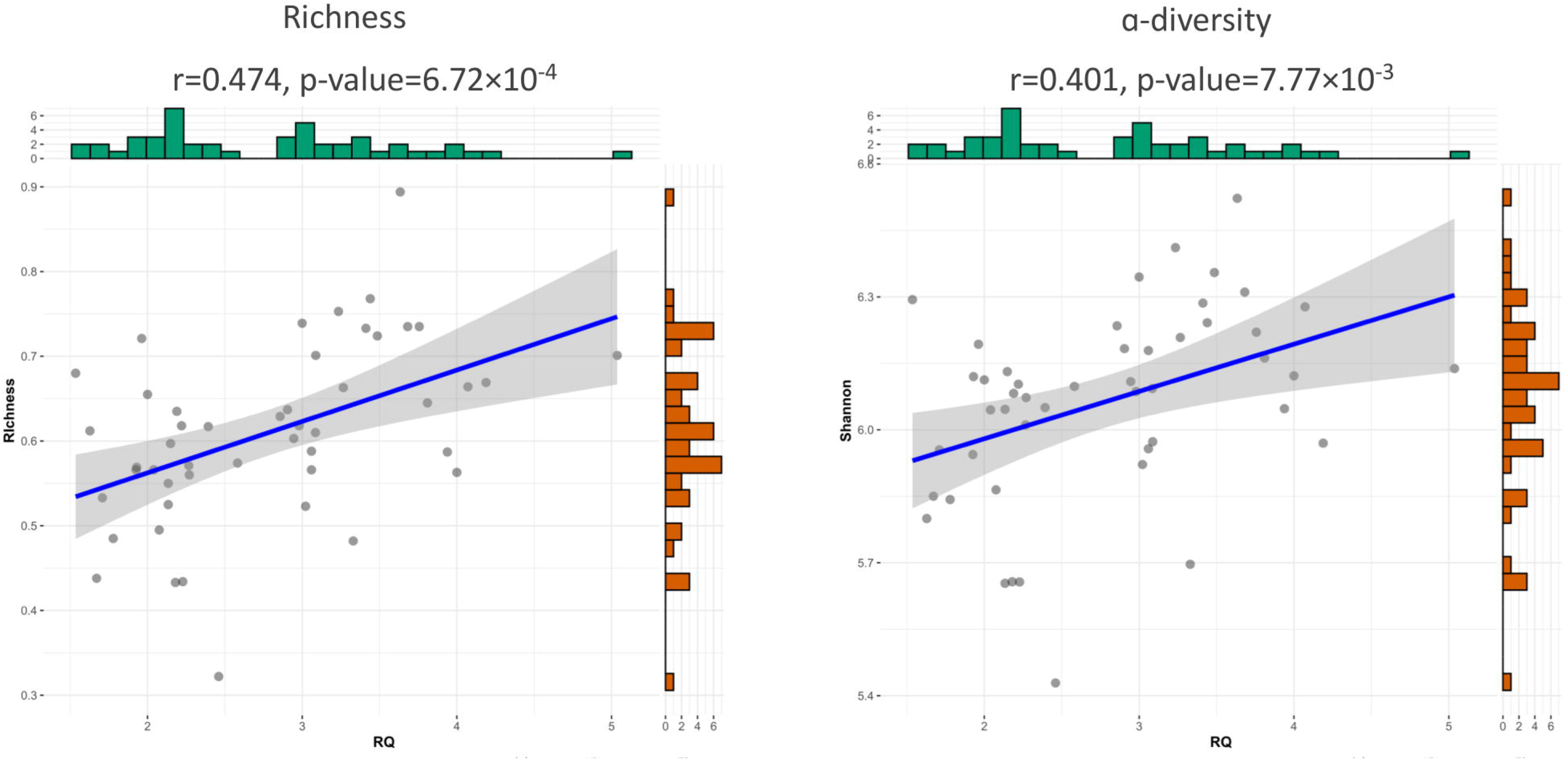
Relationship between the CNV relative quantification of the number of copies (RQ) with the richness and Shannon α-diversity index.

Remarkably, variation in the number of copies of the *ABCC2-DNMBP loci* was also segregating in an unrelated commercial F1 Duroc×Iberian crossbred pigs, with 13 out of 24 pigs showing a gain of copies. Moreover, despite differences on genetic background, age, or other environmental factors such as diet of farm of origin, the association between the *ABCC2-DNMBP loci* and the gut microbial diversity was replicated in the F1 Duroc×Iberian cross (Table 1, Figure 4). Indeed, in both Duroc and F1 Duroc×Iberian cross datasets, the qPCR reaffirmed that a gain of copies of the *ABCC2-DNMBP loci* was positively associated to the richness and Shannon α-diversity of the pig gut microbiota (Table 1).

**Figure 4.**
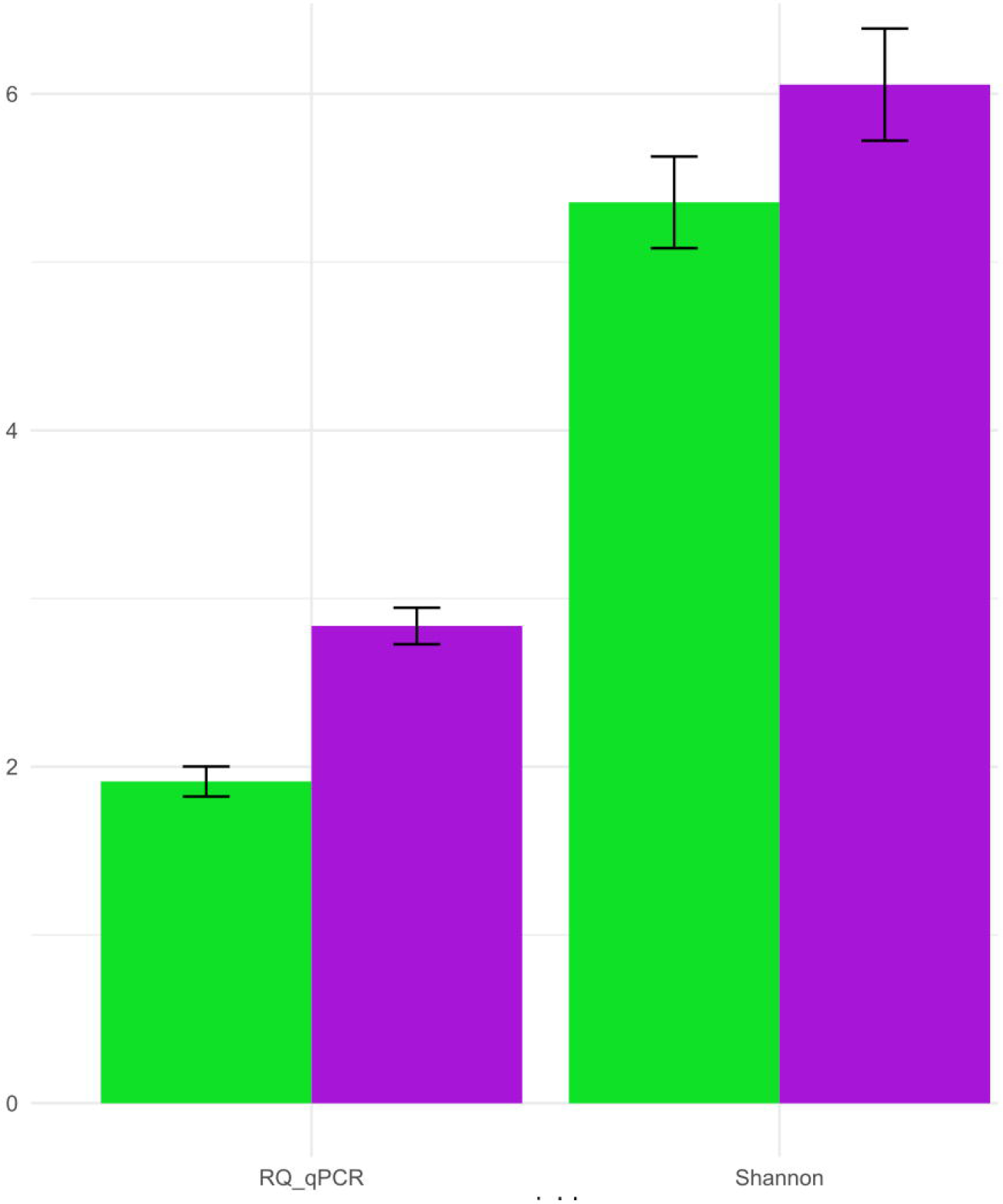
Results from the replication analysis comparing mean RQ and Shannon α-diversity in the F1 Duroc×Iberian cross. Green represents diploid (2N) samples, and purples ones DUP samples.

### Microbial signatures linked to variation in the number of copies

Results from the supervised classification model showed that the relative abundance of 122 ASVs allowed the classification between groups of DUP vs 2N samples (Supplementary table 2). The taxa-set enrichment analysis pointed out a higher overall discriminant importance of ASVs members of the *Desulfovibrio, Blautia, Phascolarctobacterium*, *Fibrobacter, Roseburia, Faecalibacterium, Megasphaera, Succinivibrio, Coprococcus, RFN20*, and *Anaerovibrio* genera (Figure 5A). Furthermore, supporting their discriminative role, we observed that compared to their diploid counterparts, the gut microbiota of DUP pigs exhibited a higher relative abundance (FDR<0.05) of the *Desulfovibrio, Blautia, Phascolarctobacterium, Faecalibacterium, Megasphaera, Succinivibrio* and *Anaerovibrio* genera, but lower relative abundance of the *Fibrobacter* and *RFN20* genera (Figure 5B). To be noted, the results obtained from the differential abundance analysis done on the unrelated F1 Duroc×Iberian crossbred population confirmed a higher relative abundance of the *Desulfovibrio, Blautia, Phascolarctobacterium, Faecalibacterium, Succinivibrio* and *Anaerovibrio* genera in the gut microbiota of samples with a gain of copies of the *ABCC2-DNMBP loci* (Figure 6).

**Figure 5.**
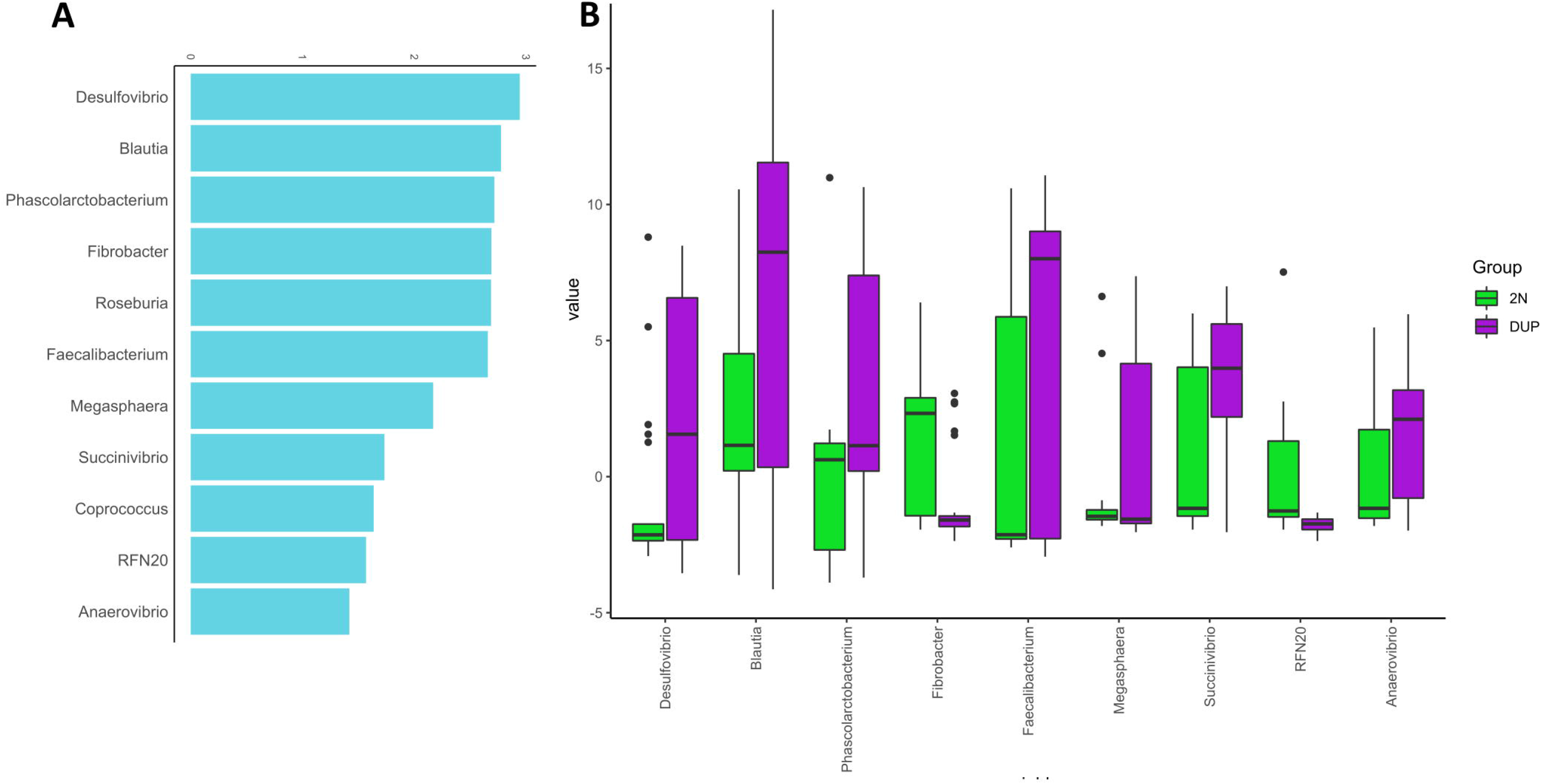
Results from microbial signature analyses at Genus level. A) Taxa-set enrichment. B) Patterns of differential abundance analysis between DUP and 2N pigs in the purebred Duroc population.

**Figure 6.**
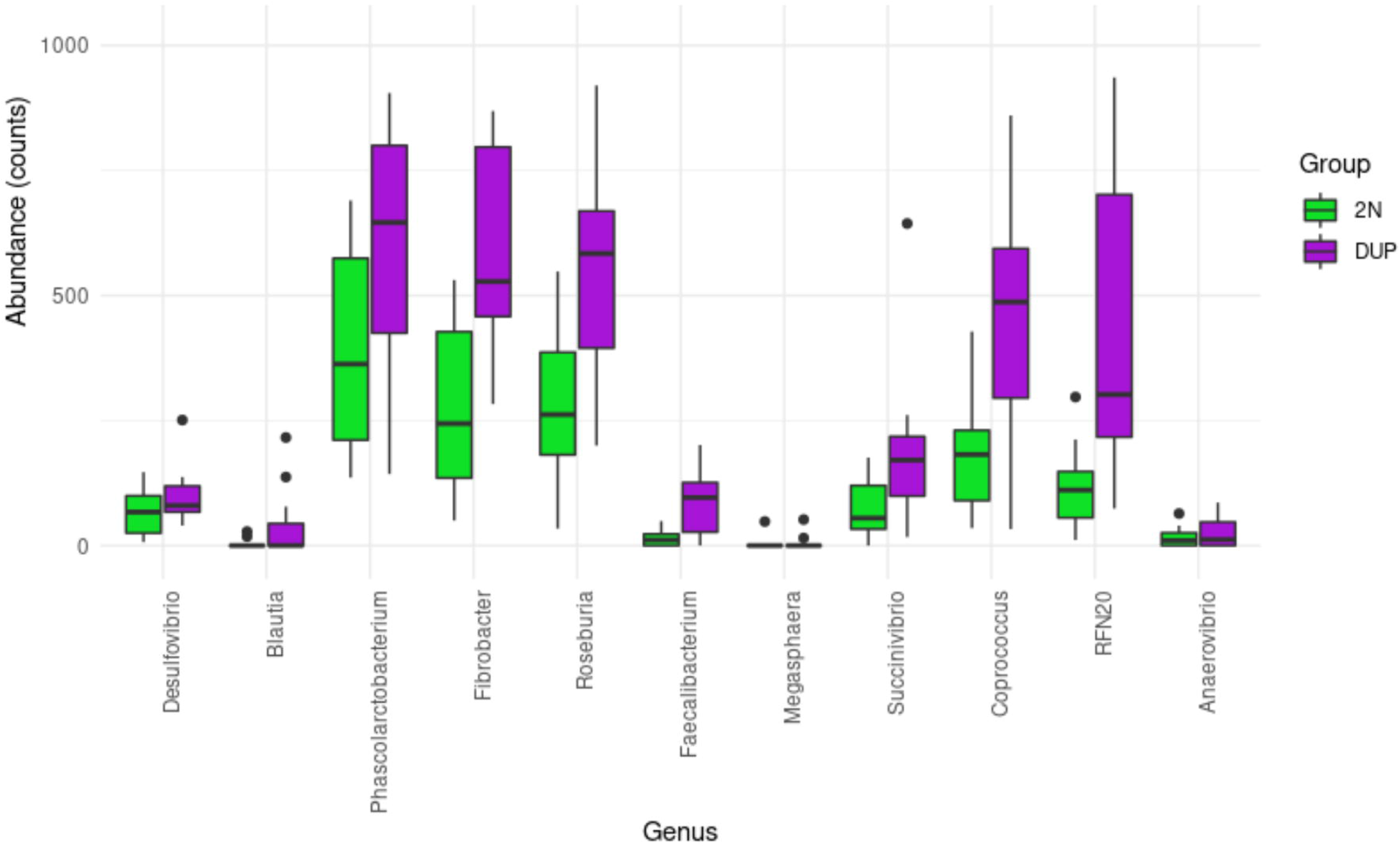
Differential abundance patterns at genus level between DUP and 2N samples in the F1 Duroc×Iberian crossbred population.

## DISCUSSION

In this study we report, for the first time in a livestock species, a CNV partially containing the *ABCC2* and *DNMBP* genes associated to the diversity and composition of the pig gut microbiota. *ABCC2* encodes a multidrug resistance-associated protein 2 (MRP2) that plays a relevant role preserving hepatic and intestinal homeostasis [32]. ABCC2 is involved in the excretion of conjugated bile acids (BAs), bilirubin, xenobiotics, and the transport of other organic anions [33][34]. In pigs, *ABCC2* has reported as co-associated to the intramuscular profile of fatty acid composition in an Iberian×Landrace cross [35]. In addition, the genomic interval harboring the *ABCC2-DNMBP loci* overlapped with QTLs associated to muscle profile of palmitic (QTLId: 95385), stearic (QTLId: 95386) and palmitoleic (QTLId: 95387) fatty acids content in a Duroc×(Landrace×Yorkshire) cross [36]. In other species such as mice, rat or humans, mutations in *ABCC2* are related to hereditary liver diseases. *Mrp2^−/-^* mice are viable [37] [38], but like *ABCC2*-knockout rats, showed chronic hyperbilirubinemia followed by a reduction in biliary excretion of bilirubin glucuronides [39] [37]. Meanwhile, mutations in the human *ABCC2* gene results in the Dubin-Johnson syndrome, an autosomal recessive disorder characterized by a defect in the transport of endogenous and exogenous anionic conjugates from hepatocytes into the bile [40]. It is worth to highlight that a genomic duplication of 5,299 base pairs comprising exons 24 and 25 of human *ABCC2* gene was predicted to result in the insertion of a premature stop codon [41].

Considering the key role of *ABCC2* on the excretion of bilirubin and conjugated BAs, we hypothesize that variation in the number of copies of *ABCC2* may influence gut levels of conjugated BAs and/or bilirubin. A bidirectional crosstalk between gut microbiota and the metabolism of bilirubin and conjugated BAs has been documented. Bilirubin can regulate the composition of gut microbiota by being potentially toxic towards Gram-positive bacteria, while promoting the proliferation of Gram-negative species [42]. In a similar way, a higher BA tolerance is evidenced by Gram-negative bacteria [43]. In agreement with these studies, the gut microbiota of DUP samples in both the discovery and validation datasets showed a higher relative abundance of Gram-negative bacteria, such as members of the *Desulfovibrio, Phascolarctobacterium, Faecalibacterium, Succinivibrio*, and *Anaerovibrio* genera. On another note, gut microbiota composition can regulate BA and bilirubin production and signaling. Evidence from germ-free (GF) rats reveals that gut microbiota is a key player in the reduction of bilirubin to urobilinoids with significant lower fecal urobilin levels in GF rats compared with conventional ones [44]. Regarding BAs, GF mice showed significant differences on enterohepatic circulation and BA composition compared with conventional mice [45] with a lower proportion of secondary and tertiary unconjugated and glycine-conjugated BA in tissues of GF rats [46]. In addition, conjugated BAs can have a protective role on gut barrier integrity [47]. The oral administration of two major conjugated BAs, tauro-cholic acid and β-tauro-murocholic acid, increased the richness of neonatal small intestinal microbiota with a positive effect on the postnatal microbiota maturation [48]. To be noted, among the top discriminant ASVs we observed butyrate producer species with a potential beneficial effect for the host, such as *Blautia obeum (ASV2433, ASV2171, ASV2278), Faecalibacterium prausnitzii (ASV2371, ASV2378), Butyricicoccus pullicaecorum (ASV2567*) and *Roseburia faecis (ASV1822*). Interestingly, the genome of all these species encodes bile salt hydrolases (*BSH*, EC 3.5.1.24) [49] [50], enzymes that mediate the primary BA deconjugation and successive conversion to secondary BAs. Therefore, partly determining the amount of secondary BAs in the colonic epithelium, which in turn acts as signaling molecules mediating different metabolic processes interconnected with health and diseases [51] [52].

The CNVR454 also included *DNMBP*, a gene that regulates the structure of apical junctions through F-actin organization in epithelial cells [53]. *DNMBP* is also involved in luminal morphogenesis and enterocyte polarization [54] [55], thus potentially contributing to the function and homeostasis of intestinal epithelial barrier (IEB). In fact, the crosstalk between IEB and the gut microbiota is crucial for the maintenance of intestinal homeostasis. For example, enterocytes, which are the most abundant population among intestinal epithelial cells, express a range of pattern recognition receptors for sensing the microbe-associated molecular patterns. Further, enterocyte apex is covered by thousands of microvilli that are vital in colonic wound repair and the transport of molecules and nutrients such as bile salts, electrolytes and vitamins [56] [57] [58] [59]. Interestingly, depletion of microbiota in mice resulted in altered patterns of microvilli formation [60]. Likewise, compared with conventional piglets, germ-free (GM) pigs displayed aberrant intestinal morphology with longer villi and shorter crypts. Meanwhile, the oral administration of commensal bacteria increased crypt depth, and induced enterocyte brush border microvilli enzyme activities on these GM piglets [61] [62] [63] [64]. Therefore, considering the functional roles of *DNMBP* in the IEB, we cannot rule out the contribution of *DNMBP* to the modulation of the diversity and composition of the pig gut microbiota.

Altogether, our results pinpointed a positive association of the variation in the number of copies of the *ABCC2-DNMBP loci* with the richness, α-diversity, and composition of the pig gut microbial ecosystems. Such findings open the possibility to modulate the gut microbiota, which has emerged as a promising breeding or therapeutic tool to optimize livestock production efficiency, animal health and well-being. A greater gut microbial diversity is usually desired, and generally accepted as an indicator of a resilient microbial ecosystem, gut, and host health. Indeed, a diverse and healthy gut has a positive effect on the absorption of dietary nutrients, feed efficiency and animal well-being.

We are aware of some limitations of our study like the limited taxonomic resolution achieved by targeting the V3-V4 16S rRNA genomic region with short-read sequencing. We are also awake about the convenience of performing further analyses to confirm the raised hypotheses by assessing the metabolic profile of BAs as well as evaluating the role of the CNV on gene expression (at both microbial and host-level) of genes involved in BA metabolism. Despite these limitations, our findings contribute to the understanding of host-microbiome interactions. Moreover, our results open the possibility to breed the holobiont via the incorporation of this source of variation on custom-made arrays that can be used in routine genotyping tasks applied to breeding programs, and together with nutritional or management strategies, will favor the simultaneous improvement of microbial traits, gut health, and host-performance.

## CONCLUSIONS

Here we report the first study exploring associations between porcine CNV and the diversity and composition of the pig gut microbiota. We identified, functionally validated, and replicated in an unrelated population a positive association between the gain of copies of *ABCC2-DNMBP loci* and the composition and diversity of the pig gut microbiota. These results suggest a role of the host-genome structural variants in the modulation of microbial ecosystems, and open the possibility of including CNVs in selection programs to simultaneously improve microbial traits, gut health and host-performance.

## Supporting information

Description of the 531 Copy Number Variant Regions

Taxonomic composition at family level of the 122 discriminant ASVs

Correlation between CNV relative quantification (RQ) and the nucleotide variability of the CNV genomic interval estimators ATajima and RTajima

## Supplementary Data

**Table S1.** Description of the 531 Copy Number Variant Regions.

**Figure S1.** Correlation coefficients between CNV relative quantification (RQ) and the nucleotide variability of the CNV genomic interval estimators ATajima and RTajima.

**Table S2.** Taxonomic composition at family level of the 122 discriminant ASVs.

## ABBREVIATIONS

QIIME2: Quantitative insights into microbial ecology
GWAS: Genome-wide association studies
SNP: Single nucleotide polymorphisms
CNV: Copy number variants
CNVR: Copy number variant region
DUP: CNV predicted as gain
BA: Bile acids
ATajima: Tajima’s theta estimator or nucleotide diversity
RTajima: Relative Tajima’s theta estimator

## DECLARATIONS

### Ethics approval and consent to participate

Animal care and experimental procedures were carried out following the institutional guidelines for the Good Experimental Practices and the Spanish Policy for Animal Protection RD 53/2013, which meets the European Union Directive 2010/63/EU for protection of animals used in experimentation, and were approved by the IRTA Ethical Committee. Consent to participate is not applicable in this study.

### Consent for publication

Not applicable.

### Availability of supporting data

The raw sequencing data employed in this article has been submitted to the NCBI’s sequence read archive (https://www.ncbi.nlm.nih.gov/sra); BioProject: PRJNA608629.

### Author Contributions

YRC, DCP and MB designed the study. OGP and MB carried out the DNA extractions. YRC, MB, OGP and RQ, performed the sampling. YRC, DCP, MB, JM, CS, SRO, and KGA analyzed the data. YRC, DCP, RQ, JMF, SRO and MB interpreted the results and wrote the manuscript. All the authors read and approved the final version of the manuscript.

## Acknowledgements

The authors warmly thank all technical staff from *Selección Batallé* S.A for providing the animal material and their collaboration during the sampling.

