## Supplementary figures and images for "Copy number variation on *ABCC2-DNMBP loci* impacts the diversity and composition of the gut microbiota in pigs"

### Correlation between CNV relative quantification (RQ) and the nucleotide variability of the CNV genomic interval estimators ATajima and RTajima

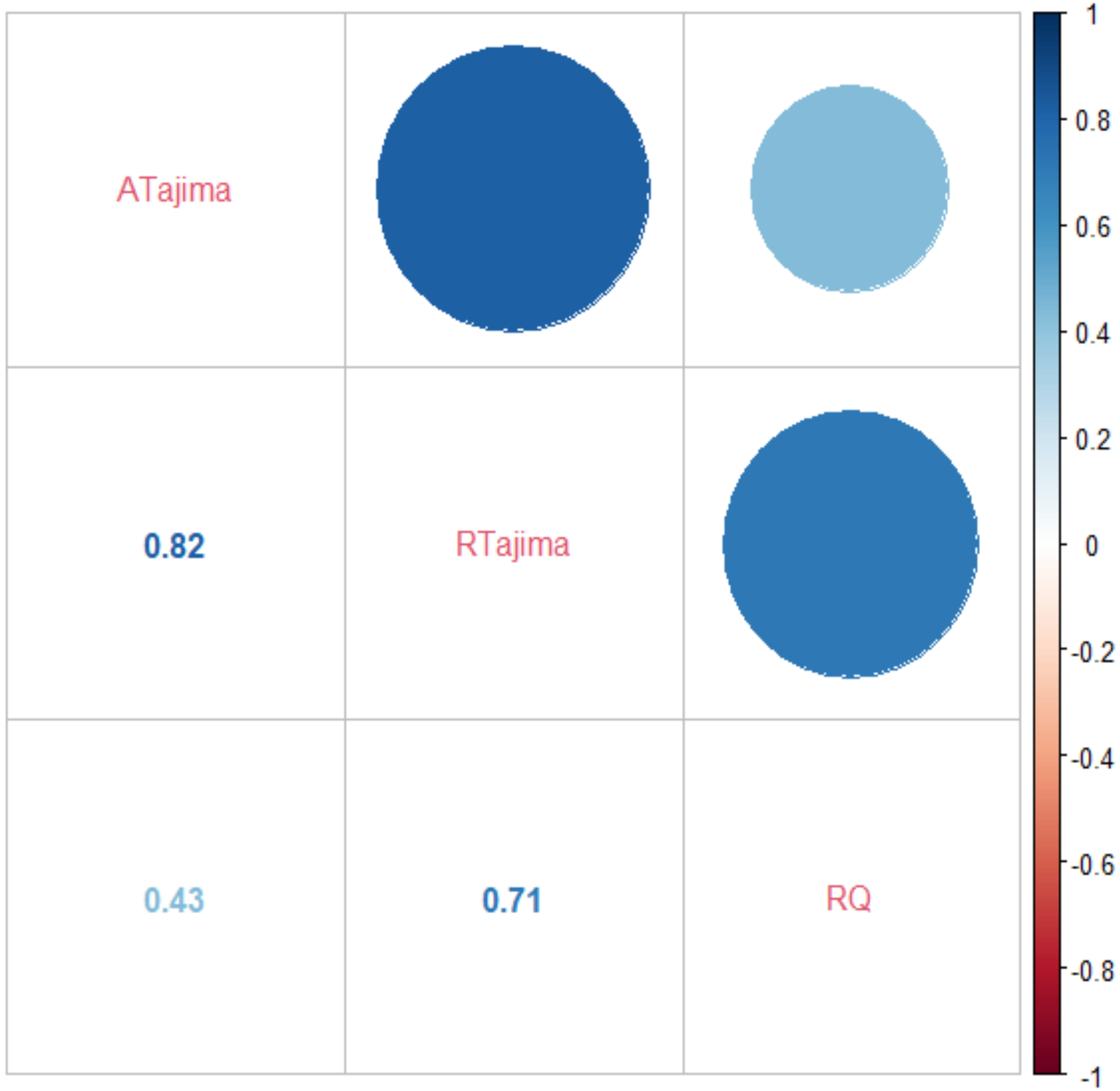
